# From Field Photosynthesis to Genetic Architecture: Insights from the First Dedicated Photosynthesis Hackathon

**DOI:** 10.64898/2026.07.24.740625

**Authors:** Anna Matuszyńska, Olakunle Sansa, Femi James Adekoya, Olusegun Olaitan Akinyemi, Esther Anokye, Olukunle Babatunde Bashir, Zsa Zsa Friederique Boyny, Mkpuma Kenneth Chukwuka, Elouen Corvest, Adenike Oluwaseun Dada, Matteo Dell’Acqua, Georgina Lala Ehemba, Andris Justin Finkbeiner, Swivia Hamabwe, Djidjoho A.T. Hodehou, Olivia Kacheyo, Kelvin Kamfwa, Kanthu Joseph Mhango, Wajiha Mu’az Abdullahi, Gideon Munduwe, Solomon Ntukidem, Olalekan Kazeem Obisesan, Ifeoluwa Simeon Odesina, Nwogu-Uchenna Ogechi, Omotola Dorcas Olaoye, Mary Morenike Olayinka, Isaac Osei-Bonsu, Kolawale Olalekan Rilwan, Leandro Stival, Zeynu Tehar, Regina Mumbua Tende, Jacky To, Uchendu Kelechi Ugochukwu, Anika Unger, Marvin van Aalst, Dominik Vrbic, Chao Zhang, Tom P.J.M. Theeuwen, David M. Kramer, Johannes Kromdijk

## Abstract

Photosynthesis is among the most consequential yet genetically complex traits in crop plants, and translating its natural variation into actionable genomic targets remains a central challenge for breeding climate-resilient varieties. To start addressing this, researchers are generating increasingly large, multi-environment field photosynthesis datasets. Yet, these data have been structurally under-analysed since their inception. Here we report the outcomes of the first dedicated hackathon focused on computational mining of such field data held in Accra, Ghana, in March 2026. Bringing together data scientists, plant physiologists, geneticists, and breeders from Europe and Africa, these interdisciplinary teams used photosynthetic data collected with hand-held fluorometers to genome-wide marker data across four crop species: cowpea (*Vigna unguiculata*), barley (*Hordeum vulgare*), common bean (*Phaseolus vulgaris*), and potato (*Solanum tuberosum*).

Despite using different species and methods, independent teams identified the same three key findings. First, mechanism-informed feature engineering and dynamic modelling recover genetic signals that are not detected or discarded in standard analysis pipelines, resulting in traits with improved heritability and meaningful associations with yield. Secondly, machine learning methods proved effective at uncovering genetic associations, with temporally resolved features substantially outperforming single time-point measurements. Third, raw chlorophyll fluorescence and absorbance traces consistently contained more information and predictive power than the extracted parameters currently used. A defining feature of this event was having experimentalists and data scientists working together, enabling AI approaches to be grounded in domain knowledge and biological mechanisms rather than relying on data alone.

## 1. Introduction

Photosynthesis is the primary source of organic carbon for crop biomass, and improving its efficiency in field conditions is one of the most pursued ambitions in current plant science. Theoretical analyses place the maximum solar energy conversion efficiency of C3 photosynthesis at 4.6% under natural conditions, while most crops achieve less than half this figure in the field (Monteith, 1977; Zhu et al., 2008). The gap reflects biological inefficiencies in light harvesting, energy dissipation, and carbon fixation, all of which are genetically variable, potentially heritable, and, in principle, reducible (Smith et al., 2023). Engineering demonstrations have confirmed this potential: accelerating non-photochemical quenching (NPQ) relaxation improved tobacco biomass by up to 20% in field conditions (Kromdijk et al., 2016); photorespiratory bypass increased tobacco yield by 40% under controlled conditions and other bypasses are predicted to increase photosynthetic efficiency by 27% (South et al., 2019; Smith et al., 2025); ribulose-1,5-bisphosphate carboxylase/oxygenase modifications improved carbon fixation rates (Salesse-Smith et al., 2018; Yoon et al., 2020); and manipulation of Calvin-Benson-Bassham cycle flux improved growth in multiple species (Ainsworth & Long, 2021).

Despite these advances, progress toward genetic improvement of field photosynthesis has remained limited (Kromdijk & Long, 2016; Long et al., 2025; Ort et al., 2015; Theeuwen et al., 2022). First, phenotyping photosynthesis at scale in the field is technically demanding: gas exchange measurements are slow and expensive, and earlier fluorescence-based methods required laboratory conditions incompatible with large breeding nurseries. This enabling factor has been substantially addressed through the development of affordable handheld devices for measuring photosynthetic traits. Low-cost deployment of sufficient parallel devices of these handheld sensors has made high-throughput assessment under field conditions achievable (provided an equal number of operators can also be arranged). The MultispeQ device (Kuhlgert et al., 2016) delivers leaf-level measurements of, among others, chlorophyll fluorescence parameters (PSII quantum yield (Φ_II_), NPQ quantum yield (Φ_NPQ_), non-regulated energy loss quantum yield (Φ_NO_), fraction of open PSII centers (qL), and linear electron flow (LEF)) (Kramer et al., 2004; Tietz et al., 2017), relative chlorophyll content (SPAD), ATP synthase conductivity, and proton motive force (ECSt, vH+, gH+) (Avenson et al., 2005; Kanazawa & Kramer, 2002), PSI activity (active centers, open/oxidized/over-reduced states), as well as environmental parameters (Photosynthetically active radiation (PAR), temperature, and humidity) in approximately 15 seconds per measurement. Users are also able to develop their own protocols based on specific research interests by incorporating different light intensities or newly defined parameters. The short measuring time enables phenotyping campaigns at the scale required for multi-environment trials and genetic mapping studies. With the MultispeQ more than 6.2 million such measurements had been recorded by June 2026 (PhotosynQ and openJII platforms). Deployment of this infrastructure has enabled genome-wide association studies linking photosynthetic traits to genomic regions in cowpea (*Vigna unguiculata*) (Sansa et al., 2024) and maize (*Zea mays*) (Ali et al., 2025; Li et al., 2025), as well as the use of genomic prediction models of yield in barley *(Hordeum vulgare)* (Gao et al., 2026). This shows that the phenotypic, genomic, and computational resources necessary for this class of analysis are increasingly available.

However, even when field phenotypic data exist, their analysis is often constrained by workflows developed for simpler, static quantitative traits and the measurements may contain more informative but unexplored parameters. Such workflows are poorly suited to photosynthetic measurements, which are dynamic, environmentally sensitive, and mechanistically structured, which may, in turn, leave substantial genetic information unused. Addressing this gap requires more than applying additional statistical methods: it requires close interaction between plant physiologists, geneticists, breeders, and computational scientists to define traits that are both analytically robust and biologically meaningful. Hackathons are increasingly used as a structured format for this type of interdisciplinary problem-solving, bringing together participants from diverse backgrounds for short, intensive, goal-oriented collaborations (Chau & Gerber, 2023). They are typically time-bounded events designed to accelerate the development of ideas, prototypes, and even scientific insights, while simultaneously fostering collaboration and community-building across disciplinary boundaries. In educational and research contexts, hackathons have been shown to promote experiential learning, multidisciplinary teamwork, and problem-solving under realistic constraints, enabling participants to integrate domain knowledge with data-driven and computational approaches (Sotaquirá-Gutiérrez et al., 2025). As highlighted in our previous work (Van Aalst et al., 2025), although most empirical evidence to date stems from broader STEM contexts, the hackathon format is particularly well suited to life sciences, where fragmented expertise and heterogeneous data necessitate tightly integrated, short-cycle collaboration.

To address this gap and to benchmark competing analytical approaches on identical datasets, we organised the first hackathon dedicated entirely to the photosynthesis–genomics intersection. Held in Accra, Ghana, in March 2026, the event brought together 35 participants from various fields. Five independent teams applied distinct computational strategies to identical curated datasets for cowpea, barley, common bean, and potato, generating convergent and complementary insights that would be difficult to obtain within a single research group (summarised in Fig. 1). This paper reports the scientific outputs of that event.

**Fig. 1.**
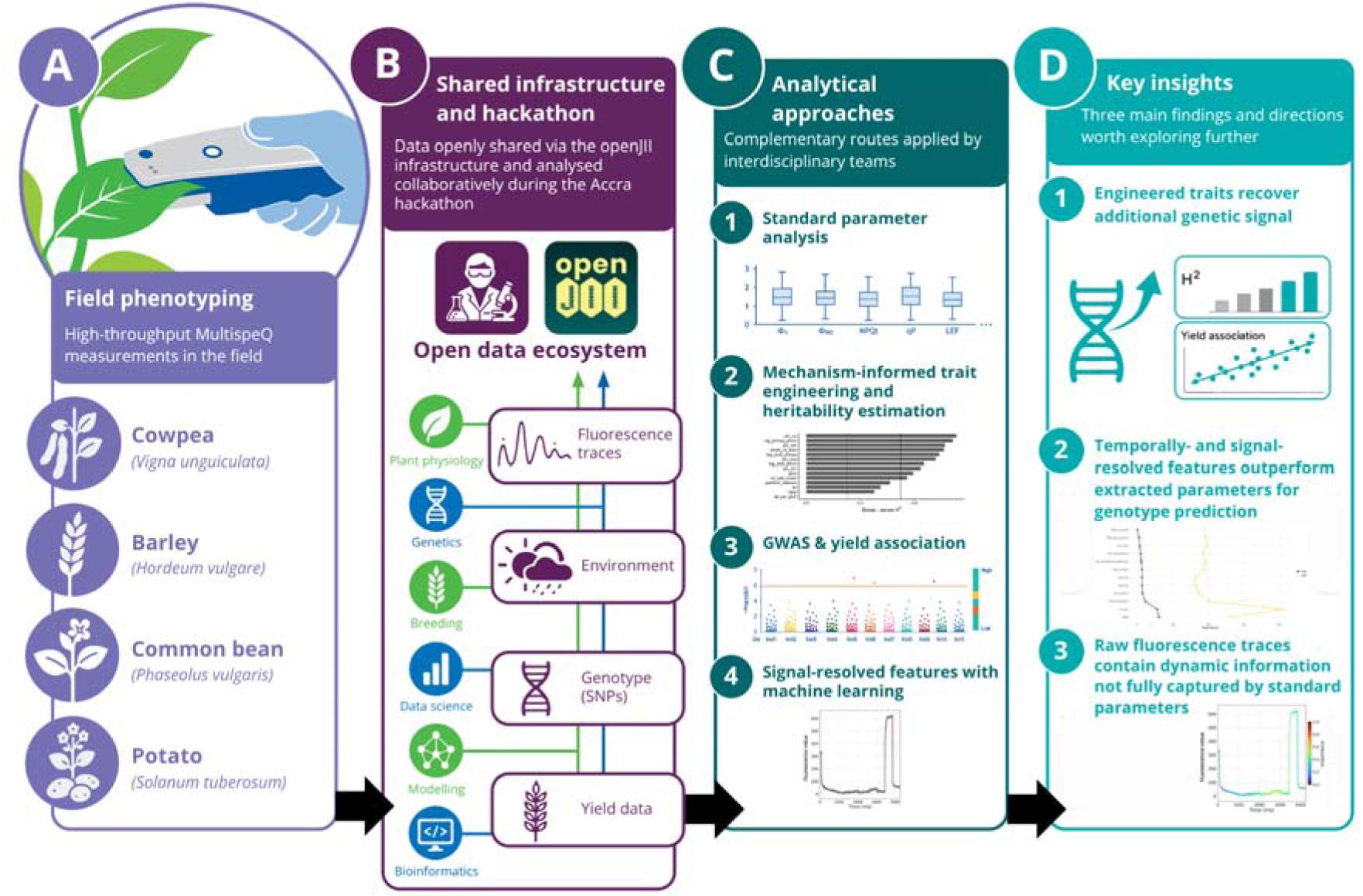
Workflow of the first dedicated photosynthesis hackathon linking field phenotyping, shared infrastructure, and genetic interpretation. Field MultispeQ measurements collected from cowpea, barley, common bean and potato included data on extracted photosynthetic parameters, raw fluorescence traces, and environmental covariates (A). They were complemented with the yield records and genome-wide marker data. During the Accra hackathon, these data layers were made available through the openJII infrastructure and analysed by interdisciplinary teams combining plant physiology, genetics, breeding, agronomy, data science, modelling and bioinformatics expertise (B). The analytical routes included standard parameter analysis, mechanism-informed trait engineering, stress-response differentials, raw-trace analysis, heritability estimation, genome-wide association studies (GWAS), yield association and machine-learning-based classification (C). The resulting insights highlight three main directions for field photosynthesis genetics: engineered traits can recover additional genetic signals, temporally- and signal-resolved features outperform extracted parameters for genotype predictions and raw fluorescence traces contain dynamic information not fully captured by standard extracted parameters (D).

Across independent analytical streams, convergent findings emerged around three themes: the heritability gains achievable through mechanism-informed feature engineering; the underexploited information content of raw fluorescence and absorbance trace data relative to already extracted parameters; and the power of machine learning to recover genetic signal when grounded in biological domain knowledge. These results are reported here as a community resource; detailed analytical treatments will follow in a series of original research articles.

## 2. The Photosynthesis Hackathon: Design and Scientific Challenge

The hackathon posed a single, well-defined challenge: identify photosynthetic parameters that reliably predict genetic components by using field-measured MultispeQ photosynthetic phenotypes paired with recorded ambient environmental data (e.g., PAR, air temperature, relative humidity) and genome-wide marker data. Teams worked across four crop species chosen to maximise biological and methodological breadth: cowpea (*Vigna unguiculata*), a drought-tolerant legume central to food security in sub-Saharan Africa (Silva et al., 2018); barley (*Hordeum vulgare*), a major temperate cereal which is often regarded as a model for cereals especially its close hexaploid relative wheat (Langridge, 2018); potato (*Solanum tuberosum*), a major staple crop with belowground storage organs (Lutaladio & Castaldi, 2009); and common bean (*Phaseolus vulgaris*) a nutritionally important grain legume and a dietary staple for millions of people (Broughton et al., 2003). All datasets were curated and delivered at the start of the event via openJII (an open-source IoT platform for recording and analysing large photosynthesis datasets, developed by the Jan IngenHousz Institute, www.openjii.org).

The programme ran for five days in Accra, Ghana, hosted by the International Institute of Tropical Agriculture (IITA): Day 1 established a shared biological and statistical foundation across disciplinary backgrounds; Days 2-4 were devoted to intensive team analysis; and Day 5 culminated in group presentations followed by planning for publication, of which this paper is the first output. Participants were 35 researchers at Master’s level and above from 14 countries across Europe and Africa, spanning data science, plant physiology, genetics, breeding, bioinformatics, and science policy. Selection prioritised geographic and disciplinary diversity, with particular encouragement for African institutions; with all participants becoming authors of this paper.

## 3. Datasets and Shared Analytical Infrastructure

All measurements were collected using MultispeQ v2 devices (PhotosynQ Inc., East Lansing, MI, USA), and experiments were designed using the PhotosynQ platform by Kuhlgert *et al*. (2016). Two measurement protocols were used depending on the dataset (see Fig. 2): the *Photosynthesis RIDES 2.0 protocol* (developed by David M. Kramer laboratory at the MSU-DOE Plant Research Laboratory, Michigan State University (MSU)), an often used protocol for MultispeQ v2 instruments running firmware ≥2.34, and the *UNZA_PIRK_DIRK_LightPotential_14 protocol,*based on the light potential protocol developed by Kanazawa *et al*. (Kanazawa et al., 2021), which is a more elaborate protocol developed by the Jan IngenHousz Institute for usage at the 2025 University of Zambia and Jan IngenHousz Institute field trials.

**Fig. 2.**
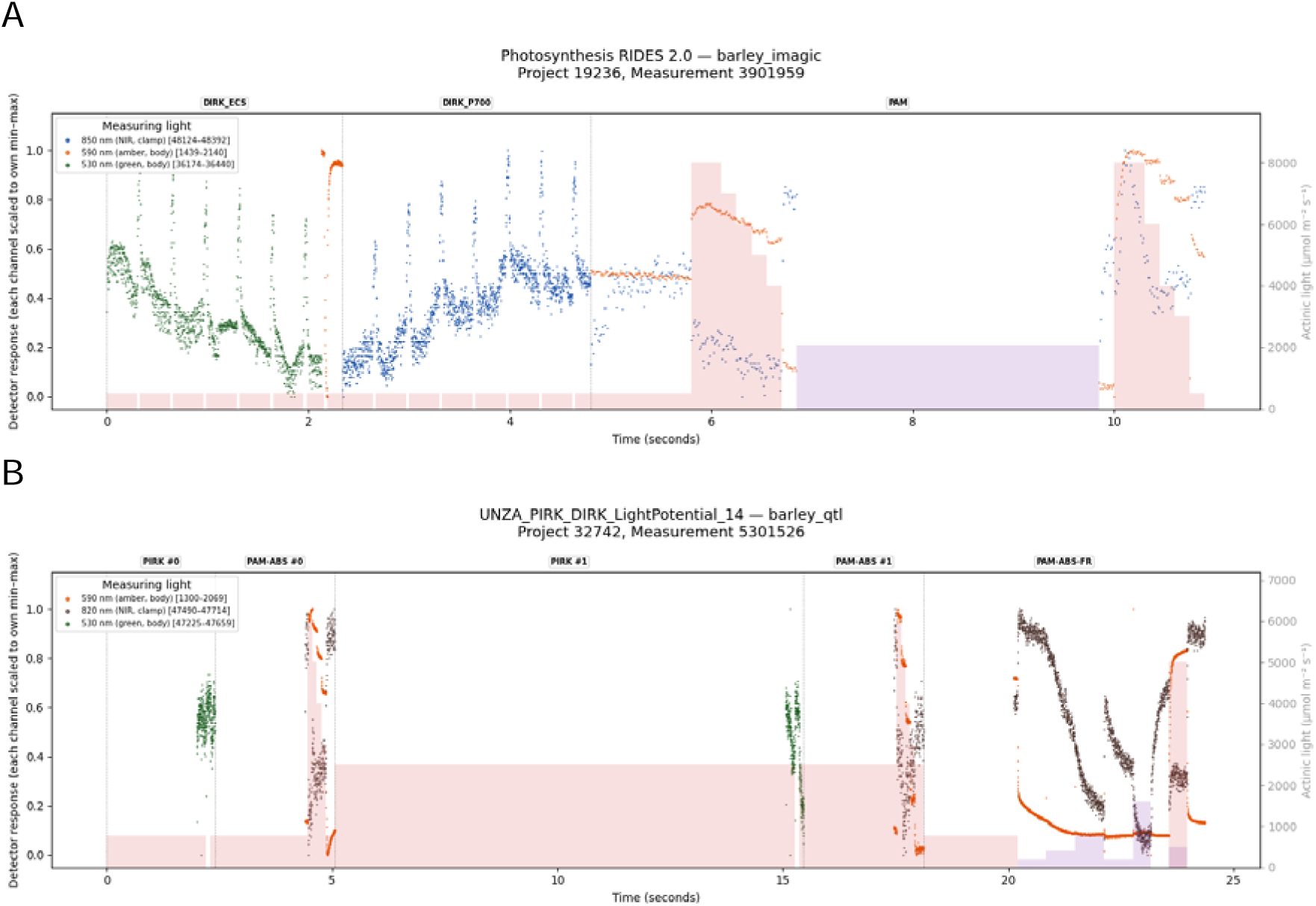
MultispeQ measurement protocols used for the field photosynthesis phenotyping datasets used during the Hackathon. Schematic representation of the A) *Photosynthesis RIDES 2.0* and B) *UNZA_PIRK_DIRK_LightPotential_14* protocols used in the analysed datasets. Both protocols record time-resolved chlorophyll fluorescence and absorbance traces together with environmental metadata, but differ in the sequence and duration of light pulses, saturating flashes and relaxation phases. Blocks of red (650nm) and purple (730nm) represent measuring light intensity and the dots represent the traces measured in green (530nm), amber (590nm) and NIR (820/850nm). Plots generated on the openJII platform (<u>openJII.org</u>).

### 3.1 Protocols

The *Photosynthesis RIDES 2.0* protocol delivers a complete measurement in approximately 15 seconds per leaf without requiring pre-acclimation by capturing ambient PAR and replicating it inside the leaf chamber using calibrated internal actinic LEDs, a key design feature that enables rapid sequential sampling across large field populations (Kuhlgert et al., 2016). The measurement sequence uses pulsed orange excitation light (605 nm, 10 µs pulses at 100 Hz), a multiphase saturating flash via the 650 nm red LED (maximum intensity 8,000 µmol photons m□² s□¹), and a farred illumination step (730 nm), producing a time-resolved fluorescence trace from which Φ_II_, Φ_NPQ_, Φ_NO_, NPQ_t_, and LEF are extracted, alongside electrochromic shift (ECS) signals recorded via dark-interval relaxation kinetics (DIRK) that estimate thylakoid proton motive force and ATP synthase conductance (Baker et al., 2007). The SPAD is recorded simultaneously via multi-wavelength absorbance. *Photosynthesis RIDES 2.0* has been validated and used for large-scale field phenotyping across multiple crop species, including common bean (Keller et al., 2024) and cowpea (Sansa et al., 2024). Throughout each measurement, ambient PAR, air temperature, relative humidity, leaf angle, and GPS coordinates are recorded automatically and stored with each observation, providing the environmental covariate dataset used in downstream modelling.

The *UNZA_PIRK_DIRK_LightPotential_14* protocol is an extension of the *Photosynthesis RIDES 2.0* protocol, adding additional light-response and relaxation blocks. Also, the multiphase saturating flash had a maximum of 6,000 µmol photons m□² s□¹. The initial actinic light period reproduces the ambient PAR, while the second actinic light period is fixed at 2500 µmol photons m□² s□¹. This allows assessment of how quickly chlorophyll fluorescence parameters respond to a sudden change from ambient to 2500 µmol photons m□² s□¹. During the far-red illumination, different light intensities are used, while chlorophyll fluorescence (excitation at 580nm, measurement at wavelengths > 730 nm) and absorbance (at about 820nm) traces are recorded. These additions extend the measurement from a rapid snapshot lasting approximately 15 s to a more detailed dynamic protocol that lasts approximately 25 s, which trades throughput for richer information and the short-term response of photosynthetic regulation to changes to high light conditions (see Fig. 2).

### 3.2 Data Sets

#### Cowpea (Vigna unguiculata), Nigeria

The MultispeQ measurements collected from a cowpea diversity panel were evaluated across multiple environments in Nigeria using the *Photosynthesis RIDES 2.0,* as described by Sansa et al. (2024). This panel provides matched pairs of before- and during-stress measurements of two years and three locations, enabling the derivation of differential traits that capture stress-response dynamics independently of baseline values. Yield data of the 112 accessions were available as a separate file and linked to photosynthetic observations via genotype × environment identifiers. DArT-Seq single nucleotide polymorphism (SNP) markers were quality-filtered as part of the published study, and participants worked only with this filtered SNP dataset for genome-wide association studies (GWAS), consisting of 9210 markers distributed across the eleven chromosome of cowpea.The final dataset for GWAS analysis of photosynthetic traits comprised ninety-eight cowpea accessions evaluated across 10 genotype × environment combinations defined by location, year, and stress conditions.

#### Barley (Hordeum vulgare), Netherlands and Ethiopia

Two independent barley datasets were analysed. The dataset from the Netherlands was collected on three recombinant inbred line (RIL) populations, which are a subset of a multiparent advanced generation inter-cross (MAGIC) population (Casale et al., 2022). The populations consisted of 272 genotypes (3 parental lines, 267 progenies, and 2 elite genotypes used as repeated checks) and phenotyped using the *UNZA_PIRK_DIRK_LightPotential_14* protocol (see Fig. 2B). The Ethiopian dataset comprised measurements of 508 genotypes from a multiparental population from three environments across two locations (Wollo and Gondar), differing substantially in altitude and ambient pressure. The data was collected using the *Photosynthesis_RIDES_2.0 protocol*. Both datasets included SNP marker data for association analysis.

#### Potato (Solanum tuberosum), Netherlands

The potato dataset was collected at eight time points within a single growing season in Wageningen, The Netherlands, providing a time-series structure absent from the other datasets. For this experiment, the *UNZA_PIRK_DIRK_LightPotential_14* protocol was used. A diverse population of forty-eight diploid potato genotypes was measured, enabling within-season trajectory analysis as well as static summaries. Due to single-season and single-location design, this dataset was used primarily for methodological demonstration of time-series approaches rather than definitive genetic mapping.

#### Common bean (Phaseolus vulgaris), Zambia

The common bean dataset focussed the photosynthetic behavior during drought stress. Four RIL populations with a total of 484 genotypes were grown in Chisamba, Zambia, during the dry season. The whole trial was irrigated, but at grain filling one half of the trial got progressively less irrigated, until no irrigation at all. For 8 days during this drought stress, MultispeQ measurements were done using the *UNZA_PIRK_DIRK_LightPotential_14* protocol.

## 4. Analytical Strategies Developed During the Hackathon

Five independent teams applied distinct but complementary analytical strategies to identical curated datasets, enabling cross-validation of findings that no single approach could have produced alone. Heritability and yield-association analyses (Section 5.1) combined linear mixed-effects models — with environment, replicate, and genotype × environment interaction as random effects, and ambient PAR, air temperature, and relative humidity as fixed-effect covariates — with genome-wide association (GWAS) analysis using Bayesian multi-locus methods that control for population stratification and kinship (Huang et al., 2019). A central methodological contribution was the treatment of PSII energy partitioning components (Φ_II_, Φ_NPQ,_ and Φ_NO_) as compositional data: because these three quantum yields sum to a fixed total, standard analysis of their absolute values can obscure genuine differences in energy allocation strategy. Isometric log-ratio (ILR) transformation was therefore applied prior to genetic analysis, alongside derivation of stress-response differential traits from paired before-stress and during-stress observations. Genotype-prediction analyses (Section 5.2) used random forest classifiers (Breiman, 2001) applied to both single time-point and temporally aggregated (AUC-transformed) photosynthetic traits. Raw-trace classification analyses (Section 5.3) used numerous machine learning approaches ranging from gradient-boosted classifiers and multi-layer perceptrons trained on extracted parameter sets to time-series foundation models applied directly to raw fluorescence traces.

## 5. Preliminary Results and Findings

Three themes structured the analyses and are reported here as the central outputs of this community effort. Detailed analyses, extended methods, and species-specific results will be reported in a series of follow-up papers currently in preparation.

### 5.1 Mechanism-informed feature engineering recovers heritable genetic signal

In both barley datasets, heritabilities for most isolated traits were near zero before environmental correction, consistent with strong masking by within- and between-day PAR, temperature, and humidity variation. After incorporating these environmental factors as fixed-effect covariates, heritabilities went up, implying a stronger recovered genetic signal. This confirms that environmental correction is not optional in field fluorescence phenotyping, it is a prerequisite for genetic signal recovery.

Analysis of the cowpea dataset showed that engineered traits designed to represent the biological mechanisms governing fluorescence signals were strongly enriched among the highest-ranked traits for broad-sense heritability (H²). In the cowpea diversity panel, comprising over 600 genotype × environment combinations across multiple Nigerian environments, ILR-transformed energy partitioning traits and log-ratio trade-off indices consistently occupied the top positions in the heritability ranking, alongside the extracted component Φ_NO_ which was the single most heritable trait (H² = 0.557). The extracted component NPQ_t_, the parameter most commonly reported in field photosynthesis studies, ranked substantially lower (H² ≈ 0.25 versus H² > 0.50 for the leading engineered traits) (see Fig. 3A). The most heritable traits reflected the Φ_NPQ_:Φ_NO_ trade-off (the balance between regulated photoprotection and unregulated energy loss) consistent with the interpretation that genotypes differ in energy allocation strategy rather than in the absolute magnitude of any single flux. Among LEF-derived traits, the composite index lef_per_phi2 was near zero in heritability, suggesting that this particular representation of linear electron flow does not capture heritable genetic variation in this population under these conditions; however, lef and lef_safe_index (explained in Table S1) showed low-to-moderate heritability and should not be generalised to the same conclusion.

**Fig. 3:**
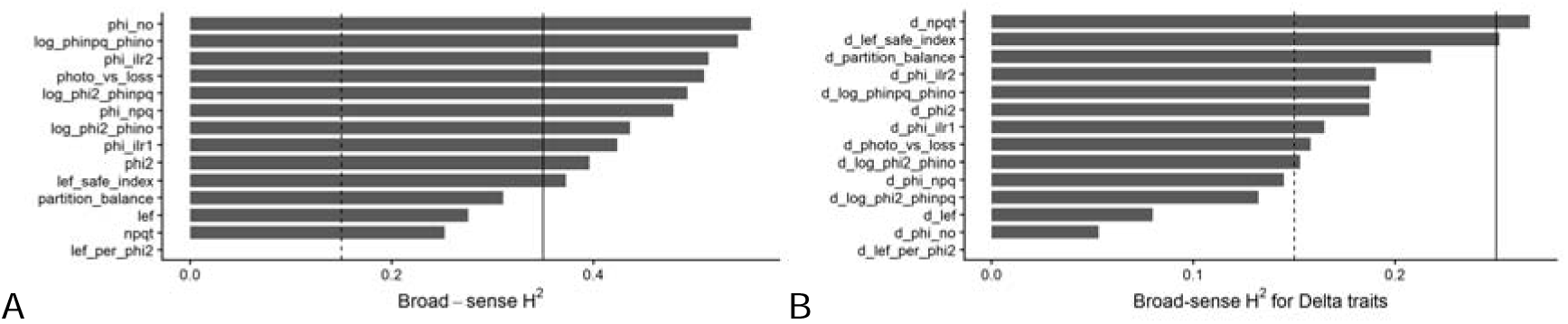
Heritability of extracted versus engineered traits. (A) Broad-sense heritability for the baseline traits as well as (B) the estimated heritability for stress-response traits. Compositional data (phi_2, phi_no, phi_npq) were transformed into ILR coordinates using an orthonormal basis defined by sequential binary partition. The first balance (ilr1) contrasts phi_2 against the geometric mean of Φ_NO_ and Φ_NPQ_ with a scaling factor of sqrt(2/3), while the second balance (ilr2) contrasts ΦNO against Φ_NPQ_ with a scaling factor of sqrt(1/2).

The paired before and during stress measurement structure of the cowpea trial enabled the derivation of fourteen stress-response differential traits. These differential traits were heritable in their own right (H² reaching 0.27 for ΔNPQ_t_), and their genetic architecture was demonstrably distinct from that of traits for either before or during stress separately, referred to as the baseline values. This can be seen in the difference between the heritability ranking across the Δ-trait panel as compared to the baseline ranking, with ΔNPQ_t_, the magnitude of enhanced non-photochemical quenching induced under stress, outranking the ILR-based differential traits that dominated the baseline panel (see Fig. 3B).

This reversal is physiologically coherent. Under ambient conditions, genotypic variation in the Φ_NPQ_:Φ_NO_ partition indicates differences in how absorbed energy is allocated between regulated photoprotective dissipation and non-regulated energy loss. Under acute stress, the primary variable distinguishing genotypes becomes whether they can activate sufficient total quenching capacity. The two trait classes therefore capture complementary aspects of photosynthetic performance that would be conflated under a single-measurement design. Genotype rankings for stress responsiveness and constitutive efficiency were not strongly correlated, providing a preliminary suggestion that breeding programmes targeting both dimensions would need to apply selection on both trait classes independently.

The engineered trade-off traits that led to the heritability ranking also showed the strongest associations with yield in the cowpea dataset. The Φ_NPQ_:Φ_NO_ log-ratio and its ILR equivalent both correlated negatively with yield (r ≈ −0.33), indicating that genotypes with a higher proportion of regulated versus unregulated dissipation tend to yield less. Stress-response traits contributed independent yield-relevant signal beyond baseline values, including delta ILR traits and ΔNPQ_t_, with fluorescence fixed effects collectively absorbing approximately 8% of genotypic yield variance in mixed-model analyses, a non-trivial proportion for a single phenotyping instrument applied in 15-second measurements per plant.

Genome-wide association analysis in cowpea recovered genomic regions exceeding the false discovery rate (FDR) significance threshold (−log₁₀(P) > 3.3) for the engineered traits d_NPQt, d_LEF Safe Index, and d_Partition Balance. While GWAS revealed significant marker–trait associations (MTAs) for these engineered traits, including some strong MTA signals at a lower significance threshold, a GWAS hit for d_npqt was identified on chromosome 7 (Vu07_4745348), corresponding to the same genomic region previously reported by Sansa *et al*. (2024) for NPQt during stress (see Fig. 4). The candidate gene (*Vigun07g046450*), located within 71 bp of the associated SNP, has functional roles in regulating plant responses to abiotic stress. The independent recovery of this locus using a different trait, here an ILR-transformed stress-response differential rather than an extracted NPQ measurement, provides biological validation that the engineered transformations preserve and in some cases amplify genuine genetic signals. Several additional candidate regions not previously reported were also identified; these likely reflect biological information specific to the relative partitioning between regulated and unregulated dissipation that is inaccessible when analysing raw NPQ values, and are prioritised in a follow-up paper focused on GWAS results.

**Fig. 4.**
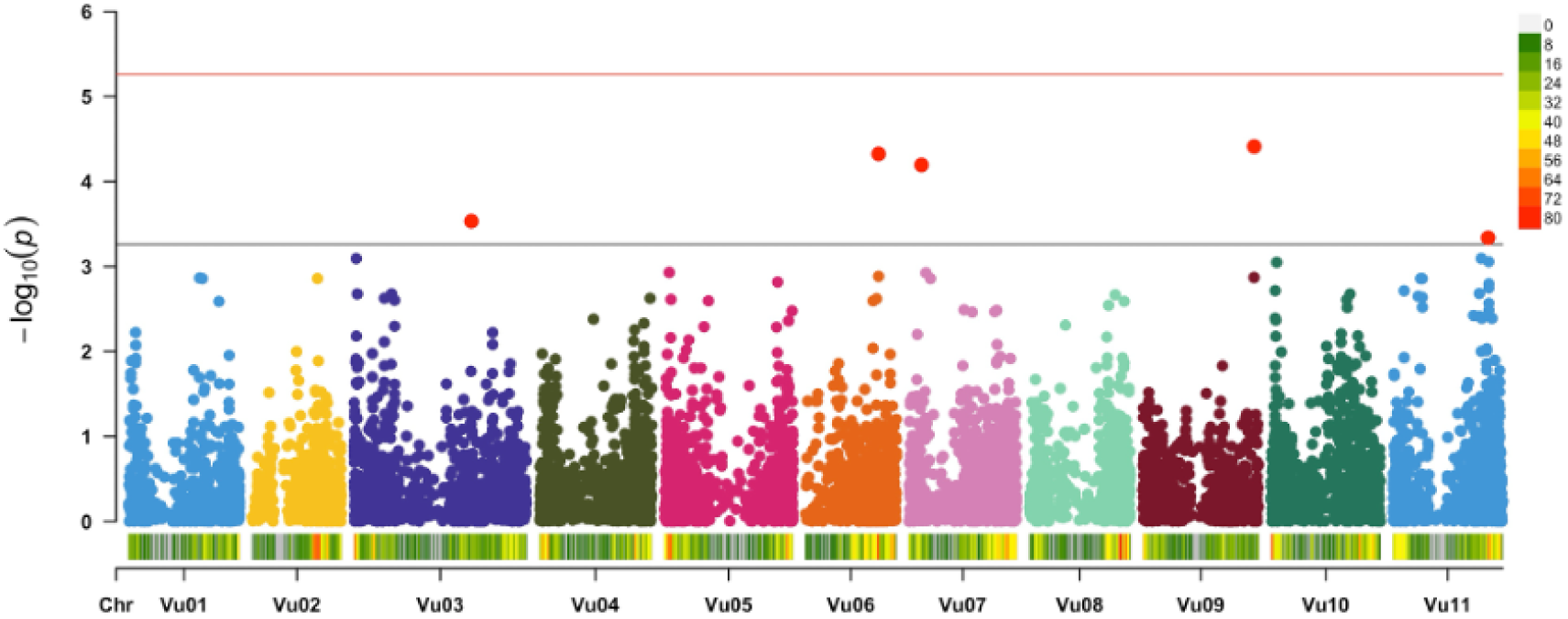
Manhattan plot showing significant MTAs for d_npqt in the cowpea dataset. 9,210 DArT-Seq SNP markers distributed across the eleven cowpea chromosomes. The red line denotes Bonferroni threshold while black line denotes the false discovery rate (FDR) threshold for declaring significant MTAs. Vu: *Vigna Unguiculata*, chr: Chromosome

### 5.2 Temporally- and signal-resolved features outperform extracted parameters for genotype prediction

Another way to assess the genetic signal is to determine whether photosynthesis-related traits contain sufficient information for genotype prediction. To this end, a random forest classifier was applied (Breiman, 2001). The random forest classifier was configured with 500 trees and validated using 100 iterations of random subsampling, with 80% of the genotypes selected for training in each iteration.

The out-of-bag (OOB) accuracy was calculated as error, where the OOB error is the proportion of incorrectly classified OOB samples.

For each tree, a bootstrap sample of the same size as the original dataset was generated by sampling with replacement. Consequently, approximately one-third of the samples were excluded from the bootstrap sample and served as OOB observations for that tree. Each sample could therefore be evaluated using only those trees for which it was not included in the corresponding bootstrap sample. The OOB error was estimated from these classifications, ensuring that each evaluated sample was classified exclusively by trees that had not been trained on it. As a result, the OOB accuracy provides an unbiased estimate of the classifier’s generalization performance.

For this approach, the potato dataset was used, which contains timeseries data on 48 genotypes (Table 1). Assuming random assignment, the probability of correctly classifying a sample by chance is 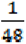. Consequently, the expected number of correct classifications under random guessing is 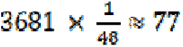 samples, that is corresponding to an expected accuracy of approximately 2.1%. When using scaled features, the mean OOB accuracy across 100 random sub-samplings reached approximately 6.7%, that is already better than random guessing.

**Table 1.**
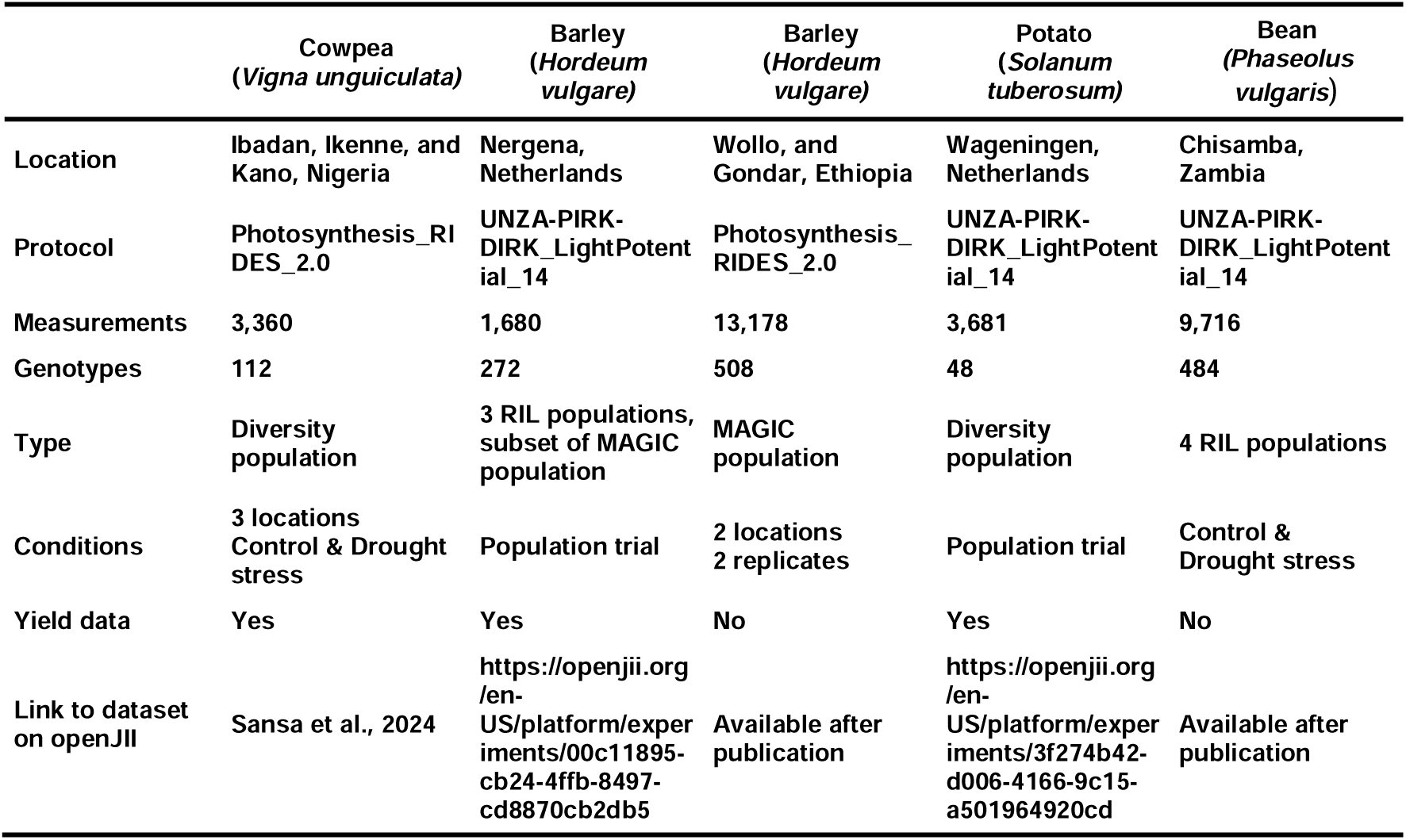
Summary of five data sets used during the 2026 photosynthesis Hackathon in Accra, Ghana.

|  | Cowpea<br>(*Vigna unguiculata*) | Barley<br>(*Hordeum vulgare*) | Barley<br>(*Hordeum vulgare*) | Potato<br>(*Solanum tuberosum*) | Bean<br>(*Phaseolus vulgaris*) |
| --- | --- | --- | --- | --- | --- |
| Location | Ibadan, Ikenne, and Kano, Nigeria | Nergena, Netherlands | Wollo, and Gondar, Ethiopia | Wageningen, Netherlands | Chisamba, Zambia |
| Protocol | Photosynthesis_RIDES_2.0 | UNZA-PIRK-DIRK_LightPotential_14 | Photosynthesis_RIDES_2.0 | UNZA-PIRK-DIRK_LightPotential_14 | UNZA-PIRK-DIRK_LightPotential_14 |
| Measurements | 3,360 | 1,680 | 13,178 | 3,681 | 9,716 |
| Genotypes | 112 | 272 | 508 | 48 | 484 |
| Type | Diversity population | 3 RIL populations, subset of MAGIC population | MAGIC population | Diversity population | 4 RIL populations |
| Conditions | 3 locations<br>Control & Drought stress | Population trial | 2 locations<br>2 replicates | Population trial | Control & Drought stress |
| Yield data | Yes | Yes | No | Yes | No |
| Link to dataset on openJII | Sansa et al., 2024 | https://openjii.org/en-US/platform/experiments/00c11895-cb24-4ffb-8497-cd8870cb2db5 | Available after publication | https://openjii.org/en-US/platform/experiments/3f274b42-d006-4166-9c15-a501964920cd | Available after publication |

Additionally, feature importance was assessed using mean decrease in accuracy (MDA) and mean decrease in Gini impurity (MDG) (Nembrini et al., 2018). For MDA, each feature was randomly permuted in the OOB samples for each tree while all other features remained unchanged. The resulting decrease in OOB accuracy, relative to the unpermuted predictions, was calculated and averaged across all trees. Features causing larger decreases in OOB accuracy were considered more important for accurate classification.

MDG was derived from the decrease in Gini impurity during tree construction. The Gini impurity was defined as 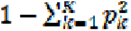, where *p_k_* is the proportion of observations in class *k*. For each split, the reduction in Gini impurity attributable to the selected feature was calculated and accumulated across all trees. MDG therefore provided an alternative measure of feature importance, with larger values indicating greater contributions to class discrimination. Across all runs, SPAD consistently emerged as the most important feature, ranking first under both MDA and MDG (see Fig. 5).

**Fig. 5:**
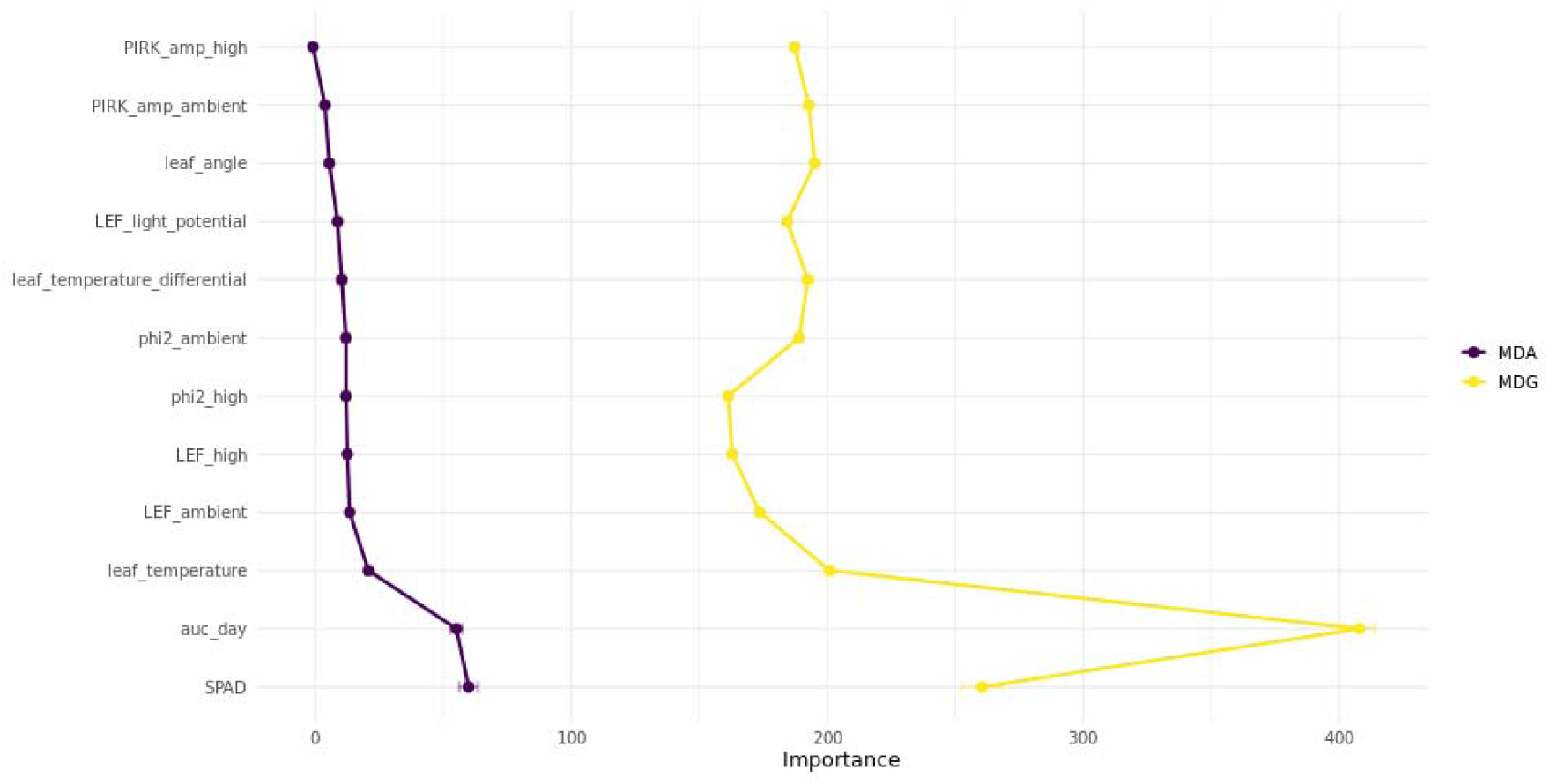
Mean Decrease Accuracy (MDA) and Mean Decrease Gini (MDG) for all features included in the random forest classifier. For Φ_II_ (phi2_high), the area-under-the-curve (AUC) was calculated separately for each day. Data from one day (2025-08-14) were excluded due to insufficient observations.

Compared to single time-point measurements of photosynthetic traits, temporally resolved dynamic features collected over the course of a measurement day appear to contain more genomic information (Kaiser et al., 2015). Ambient light, in particular, can vary substantially within minutes, leading to measurements that increasingly reflect environmental fluctuations rather than purely genotypic effects (Gao et al., 2024). Measuring all genotypes within a narrow time window captures their response to the prevailing environmental conditions at that moment, embedding the genotypic effects within that environmental context. Repeating this across multiple time windows throughout the day then allows the genotype-specific responses to continuously changing conditions to be resolved. As measuring all genotypes at precisely the same time point is not achievable in practice with handheld devices such as the MultispeQ v2, residual noise from within-window environmental fluctuations cannot be fully avoided. However, under the assumption of sufficient environmental stability within each time window, this noise may remain acceptable. Therefore, an area-under-the-curve (AUC) transformation of traits within this interval may better capture genotype-dependent photosynthetic responses. This approach was demonstrated using Φ_II_, which was measured repeatedly within a single day.

After smoothing the temporal trajectories and computing daily AUC values for Φ_II_, inclusion of these aggregated measures in the random forest model increased the OOB accuracy to approximately 28%. The AUC feature ranked highest under the MDG criterion and second highest under MDA. This indicates that time-dependent variation captures more relevant genotypic information than individual time-point measurements. However, the overall improvement in predictive performance remains limited, at least in part due to constraints in the dataset’s temporal resolution.

In addition, the substantial covariance among fluorescence-derived traits, low heritabilities, and potentially complex interactions among physiological and environmental variables suggest that relevant genotype × environment interactions may not be fully captured by individual engineered features. Future analyses may therefore benefit from integrating harmonised multi-environment datasets through shared computational infrastructures such as OpenJII. Such datasets could facilitate factor-analytic approaches that incorporate photosynthetic traits as covariates to improve the characterisation of latent structures underlying genotype × environment interactions. These frameworks represent the covariance structure of genotype × environment effects through a reduced set of latent factors, enabling efficient dimensionality reduction while capturing major interaction patterns (Kelly et al., 2007). Initial exploratory analyses from the hackathon using the Netherlands barley dataset indicated that a factor-analytical model incorporating photosynthetic covariates, including LEF light potential, PAR, SPAD, and leaf temperature differential, explained a greater proportion of variance than a baseline model. PAR emerged as the dominant source of explained variance. However, only a single latent factor could be robustly estimated due to the dataset’s environmental structure. Further evaluation across harmonised multi-environment datasets will be required to assess the broader applicability of this approach.

### 5.3 Raw fluorescence traces as a source of underused physiological information

The most striking finding across independent teams was that raw fluorescence traces retained substantially more predictive information than the extracted parameters extracted from them. In the common bean dataset, binary classifiers trained to distinguish stress-treated from control plants achieved 86% balanced accuracy using the 8 signal-separated raw traces from the *UNZA_PIRK_DIRK_LightPotential_14* protocol. For this prediction we captured performance variation across different raw trace combinations evaluated in the grid search, where *raw_pam_ambient* and *raw_pam_fr* contained most predictive power (see Fig. 6A). To see how these traces compare to the extracted parameter Φ_II_, we compared the predictive power of Φ_II_ during ambient and high light with their respective raw traces. With Φ_II_ during ambient light we reach 60% balanced accuracy, while the *raw_pam_ambient* trace reached 80% (see Fig. 6B). Combining both Φ_II_ during ambient and high light reaches 69%, which is increased to 83% when both respective traces are used. Although this result still requires validation across independent datasets, genotypes, and environments, it suggests that extracted parameters may discard dynamic information relevant to stress classification.

**Fig. 6:**
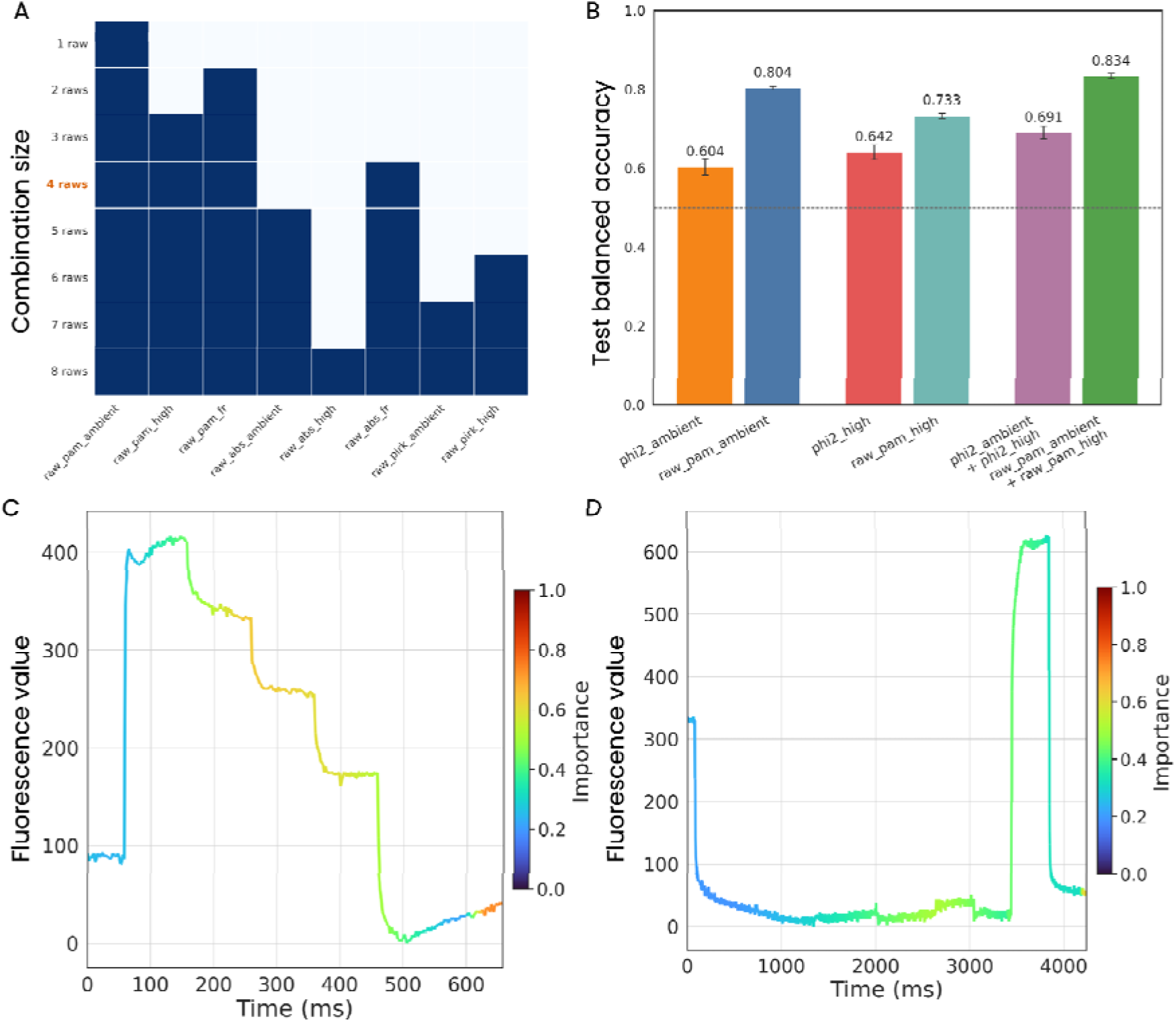
The importance of different parts of the raw fluorescence traces for binary classifiers distinguishing between stress-treated and control plants in the bean dataset. Binary classifiers were trained with all possible combinations of different sections from the traces. (A) For each total number of sections included for training (combination size), the presence of a section in the best model of that combination size is denoted by the blue rectangles. The overall best model with a classification accuracy of 86% is highlighted in orange at four sections. The importance is indicated by the frequency of a section being present in the best models, with *raw_pam_ambient* being the most frequently present. (B) The predictive power of the extracted Φ_II_ during ambient and high light as compared to their respective raw traces. (C) The feature importance of the *raw_pam_ambient* trace, which is used to calculate the extracted ΦII during ambient light. The feature importance is calculated as an average over all samples, when just this trace is used for the prediction. (D) The feature importance of the *raw_pam_fr_data* trace during the far-red associated section of the *UNZA_PIRK_DIRK_LightPotential_14* protocol. The feature importance is averaged over all samples, like in panel C.

By mapping the relative importance per trace, by averaging the importance over all samples, the middle section of the multi-phase flash pulse in the *raw_pam_ambient* trace and the rise kinetics of the saturating pulse and the response from low to high farred light in the *raw_pam_fr_data* trace contain the most information (see Fig. 2, 6C, and 6D). Its temporal structure reflects defined phases of the measurement protocol, including baseline fluorescence, light-induced changes in PSII operating efficiency, NPQ induction or relaxation, and far-red-associated dynamics. These phases are linked to processes described by dynamic models of light harvesting, photoprotection, and electron transport (Stirbet et al., 2020, 2024). The fact that machine-learning models assigned high predictive value to regions outside the standard extracted parameters, therefore, points to a biologically meaningful source of information, not merely to improved statistical classification. A promising next step is therefore to fit fluorescence traces using model-constrained learning approaches, such as physics-informed neural networks (Cuomo et al., 2022) or universal differential equations (Kidger, 2022). These methods could combine the structure of existing ordinary differential equation models with data-driven components for processes that are incompletely specified or difficult to parameterise (see Fig. 7). In practice, this would allow raw traces to be converted into model-derived traits, such as NPQ induction or relaxation rates, apparent electron transport regulation, or far-red relaxation kinetics. Such traits would be more suitable than black-box classifier outputs for heritability analysis, GWAS, and breeding applications.

**Fig. 7:**
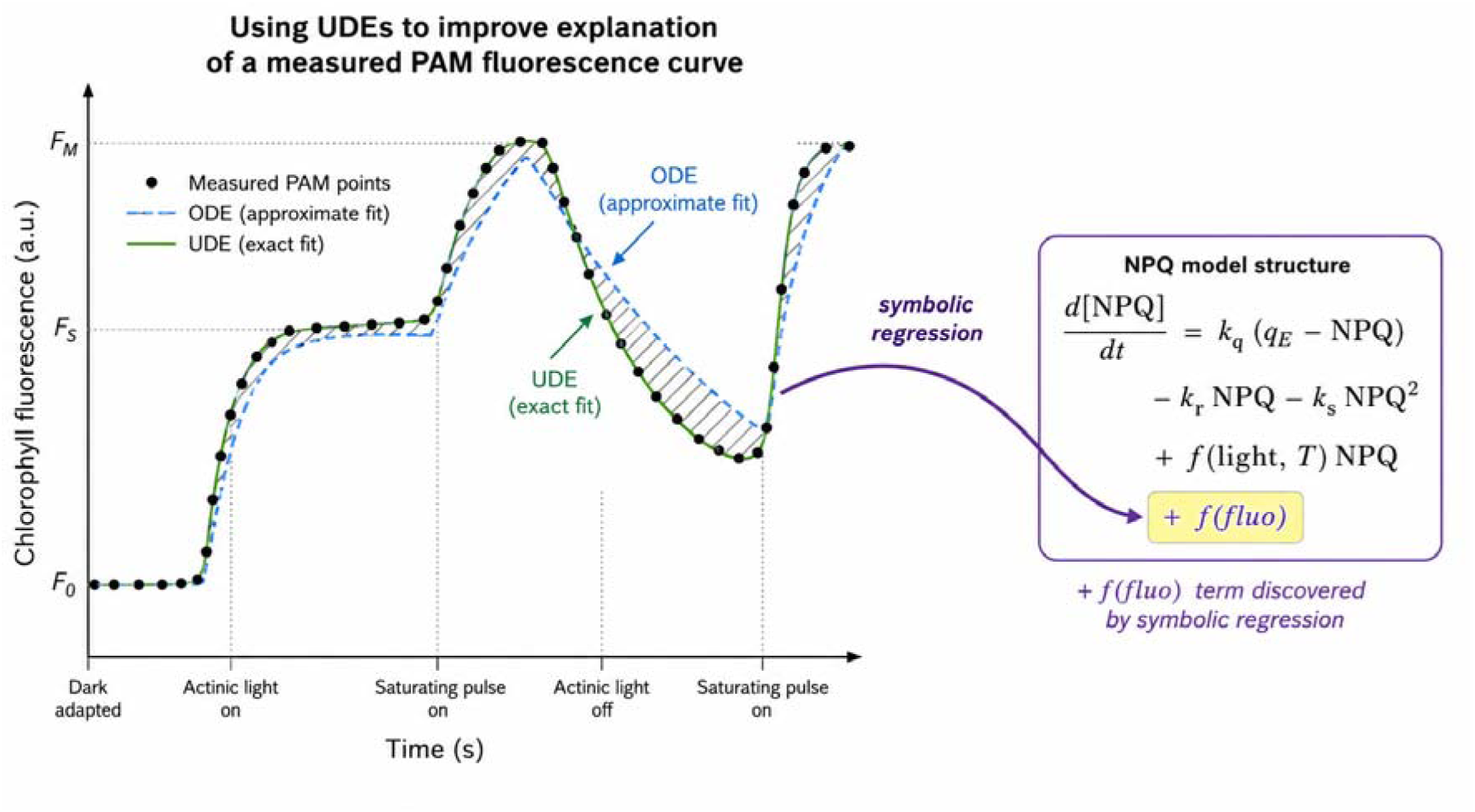
Schematic overview of using universal differential equations (UDEs) to improve the interpretation of measured PAM fluorescence traces. Measured PAM fluorescence points are first fitted with a conventional ordinary differential equation model, which captures the overall trajectory but may miss parts of the dynamic response. A UDE-based model extends the mechanistic description by learning unresolved components of the fluorescence dynamics, resulting in a closer fit to the measured trace. Symbolic regression can then be applied to the learned UDE term to recover an interpretable functional relationship, as illustrated here as an additional light-dependent term in an NPQ model structure.

## 6. Lessons for Future Photosynthesis Data Challenges

Beyond the specific biological findings, the Accra hackathon highlighted several practical requirements for productive community analysis of field photosynthesis data. First, substantial pre-hackathon data curation was essential. Phenotypic, environmental, yield, and marker datasets had to be harmonised before participants could meaningfully compare analytical strategies. In a short collaborative event, unresolved differences in file formats, genotype identifiers, environmental metadata, and trait definitions can otherwise consume the time needed for analysis.

Second, the dedicated first day was critical for establishing a shared working language. Participants differed in their familiarity with fluorescence nomenclature, MultispeQ protocols, genetic analysis, and computational workflows. It is worth noting that many participants arrived with limited prior exposure to photosynthesis as a discipline. For several, the hackathon represented their first structured engagement with fluorescence-based measurements and the underlying biophysics. Yet this was paired with a strong motivation to incorporate photosynthetic traits into their own breeding and data science work. For many, the event served as an eye-opening experience in both directions: revealing the genuine complexity of interpreting dynamic fluorescence signals, while simultaneously demonstrating that meaningful analysis is achievable within days when the right tools, data, and expertise are brought together. Short introductory sessions aligned the group on the biological meaning of the measured traits, the structure of the datasets, and the available analytical tools, allowing the following days to focus on analysis rather than disciplinary translation.

Third, shared computational infrastructure was useful, but not sufficient on its own. Providing curated datasets and analysis tools through openJII reduced installation barriers and gave participants a common starting point. However, the experience also showed that overly prescriptive environments can limit participation by requiring researchers to abandon their preferred analytical tools. Future events should therefore prioritise harmonised, analysis-ready datasets that can be easily downloaded and used in Python, R, or other environments while preserving a common data standard.

Fourth, mixed disciplinary teams shaped the analysis itself, not only the interpretation. Physiological and genetic expertise helped define biologically meaningful feature transformations, identify relevant environmental covariates, and reject statistically attractive but physiologically implausible outputs.

Computational expertise enabled participants to test approaches not yet standard in field photosynthesis genetics, including compositional transformations, time-series analysis, and machine-learning methods applied directly to raw fluorescence traces.

The event also exposed limitations. Hackathons are well-suited to generating analytical prototypes, comparing alternative approaches, and identifying cross-cutting signals, but they are not a substitute for full validation. Several findings reported here therefore require extension in companion studies with formalised pipelines, independent validation sets, and fully documented model specifications. The main value of the format was to reveal, across independent teams and datasets, where current analytical practice is likely losing information. These lessons provide a practical template for future community efforts aimed at converting high-throughput photosynthesis phenotyping into genetically and physiologically interpretable traits.

## 7. Outlook and Conclusions

The Accra hackathon showed that field photosynthesis data, already collected at scale across research projects worldwide, contain substantially more analytical potential than is captured by standard parameter-based workflows. The event brought together complementary approaches to the same problem: *how to convert dynamic, environmentally sensitive fluorescence measurements into traits that are useful for genetic analysis and breeding*. Together, our results suggest that field photosynthesis genetics should move beyond fixed parameter extraction toward trait definitions that retain dynamic, environmental, and physiological structure. Yet, these findings should be interpreted as a foundation for further work rather than as the final analysis of each dataset. The hackathon format was valuable because it allowed independent teams to compare analytical strategies quickly and generate testable hypotheses, but did not provide enough time for formal validation or provision of fully documented pipelines. Clearly, independent test sets and species-specific follow-up studies are needed. The most immediate opportunities are to standardise analysis-ready photosynthesis phenotyping datasets and to develop interpretable features from raw fluorescence traces, and make them openly available via platforms such as OpenJII.

Finally, the long-term impact of this work will depend not only on improved analytical methods, but also on expanding the community able to use them. While high-throughput field photosynthesis measurements are becoming increasingly accessible, extracting biologically meaningful information from these datasets still requires specialised expertise in plant physiology, quantitative analysis, and computational methods. Realising the full potential of these datasets will therefore require both shared expertise and effective mechanisms for transferring knowledge across disciplines and institutions. Broadening training in quantitative photosynthesis, computational analysis, and data interpretation will enable a much wider community of plant scientists and breeders to exploit these datasets, increasing the return on existing phenotyping efforts. Community teaching resources that integrate education with interactive analysis and mechanistic modelling such as the one proposed by Philipps *et al*. (Philipps et al., 2025) or Ross *et al*. (Ross et al., 2006), provide one route towards this goal, lowering the barrier to entry while promoting reproducible and physiologically informed analyses.

More broadly, this work argues for a shift in field photosynthesis genetics from extracting a fixed set of standard parameters toward designing traits that preserve physiological dynamics and environmental context. High-throughput fluorescence phenotyping can already generate the necessary data. The next advance will depend on whether these data can be converted into traits that are statistically robust, mechanistically interpretable, and useful for selection.

## Acknowledgements

The authors gratefully acknowledge funding from the Jan IngenHousz Institute, the Mastercard Foundation, University of Cambridge Climate Resilience and Sustainability Research Fund. We thank the International Institute of Tropical Agriculture (IITA) for hosting the event in Accra, Ghana, and the entire IITA local team for their generous support throughout. We acknowledge Yanrong Gao, Po-Ya Wu, Shizue Matsubara and Benjamin Stich for providing datasets that did not get used during the Hackathon. We also thank Lutz Kupferschläger (RWTH Aachen University) for the design of the overview illustration (Figure 1) for this manuscript.

## Abbreviations

AUC: area-under-the-curve
DIRK: dark-interval relaxation kinetics
ECS: electrochromic shift
FDR: false discovery rate
GWAS: Genome-wide association study
H^2^: broad-sense heritability
IITA: International Institute of Tropical Agriculture
ILR: isometric log-ratio
LEF: linear electron flow
MAGIC: Multiparent advanced generation inter-cross
MDA: mean decrease accuracy
MDG: Mean decrease gini
MTA: Marker-trait association
NPQ: non-photochemical quenching
NPQ_t_: total NPQ
OOB: out-of-bag
PAR: photosynthetically active radiation
qL: fraction of open PSII centers
RIL: Recombinant inbred line
SNP: single nucleotide polymorphism
SPAD: relative chlorophyll content
Φ_II_: PSII quantum yield
Φ_NO_: non-regulated energy loss quantum yield
Φ_NPQ_: NPQ quantum yield

